# The Modular μSiM Reconfigured: Integration of Microfluidic Capabilities to Study in vitro Barrier Tissue Models under Flow

**DOI:** 10.1101/2022.03.28.486107

**Authors:** Mehran Mansouri, Adeel Ahmed, S. Danial Ahmad, Molly C. McCloskey, Indranil M. Joshi, Thomas R. Gaborski, Richard E. Waugh, James L. McGrath, Steven W. Day, Vinay V. Abhyankar

**Affiliations:** Department of Biomedical Engineering, Rochester Institute of Technology, Rochester, NY, 14623, USA; Department of Biomedical Engineering, University of Rochester, Rochester, NY, 14627, USA

**Keywords:** tissue barrier, modular, microfluidic, microphysiological system, tissue chip

## Abstract

Microfluidic approaches to study tissue barriers have emerged to address the lack of fluid flow in conventional “open-well” Transwell™-like devices. However, microfluidic techniques have not achieved widespread usage in bioscience laboratories because they are not fully compatible with traditional, tried-and-true experimental protocols. To advance barrier tissue research, there is a need for a platform that combines the advantages of both conventional open-well and microfluidic systems. Here, we develop a plug-and-play flow module to add on-demand microfluidic capabilities to a **m**odular **micro**fluidic system featuring a **si**licon **m**embrane “m-μSiM” as an open-well device with live-cell imaging capabilities. The magnetic latching assembly of our design enables bi-directional reconfiguration between open-well and fluidic modes. This design feature allows users to conduct an experiment in an open-well format with established protocols and then add or remove microfluidic capabilities as desired. Our work also provides an experimentally-validated flow model to help select desired flow conditions based on the experimental needs. As a proof-of-concept, we demonstrate flow-induced alignment of endothelial cells and visualize different phases of neutrophil transmigration across an endothelial monolayer under flow. We anticipate that our reconfigurable design will be adopted by both engineering and bioscience laboratories due to the compatibility with standard open-well protocols and the simple flow addition capabilities.

## 1. Introduction

Endothelial-lined vascular barriers separate blood from the surrounding tissue and play a crucial role in maintaining homeostasis by regulating molecular permeability, recruiting immune cells, and preventing pathogen entry into the tissue compartment **(Figure 1A)**.^[1–3]^ Impaired barrier function or breakdown can have significant implications on human health, including stroke,^[4]^ cancer,^[5]^ cardiovascular disease,^[4,6]^ and neurodegenerative disorders.^[7,8]^ As a scalable companion to animal studies, in vitro culture models aim to replicate key aspects of the in vivo environment and are a popular method to study factors that affect barrier structure and function in a controlled manner.^[2,8–11]^

**Figure 1.**
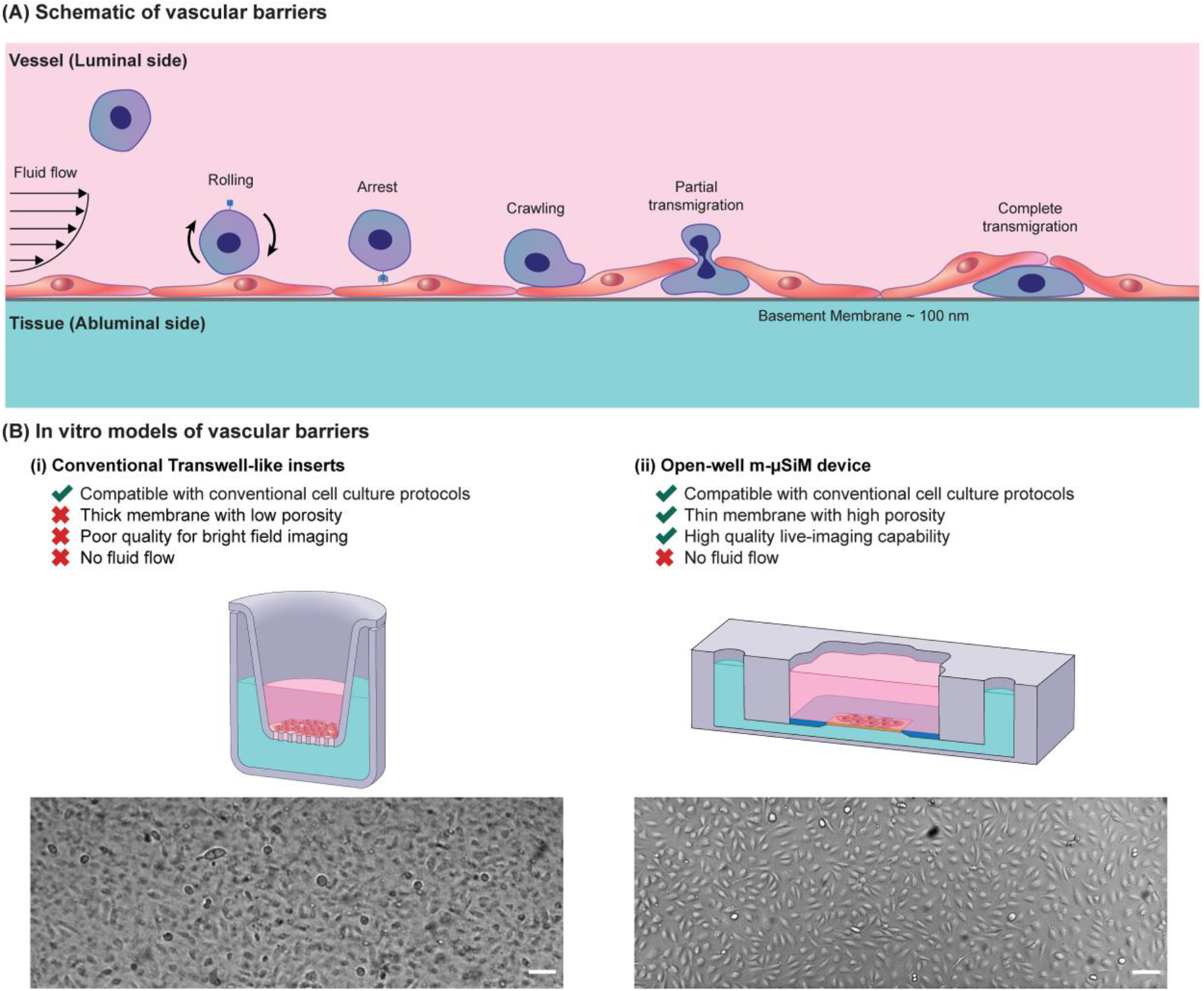
(A) Schematic illustration of an in vivo vascular barrier. The pink domain represents fluid in the luminal side, and the green domain represents the abluminal side of the barrier. Fluid flow applies shear stress to endothelial cells and carries neutrophils to the barrier. During injury, neutrophils are recruited from the bloodstream in a coordinated process of rolling, arrest, probing, and transmigration across the cell monolayer under continuous flow conditions. To mimic the vascular environment in vitro models, the membrane is used as a physical support for endothelial cells. (B) Comparison of the membrane characteristics and imaging quality between (i) conventional Transwell-like inserts and (ii) the open-well m-μSiM. Scale bars = 100 μm.

Barrier models, including the popular “open-well” Transwell™ - like format, aim to replicate the compartmentalization found in vivo by introducing a porous culture membrane separating the top and bottom media-filled compartments.^[8,11]^ Cells are seeded directly onto the membrane that physically separates the compartments while permitting soluble factor communication between them.^[8,11]^ This configuration is relevant to several barrier tissues, including the blood-brain barrier,^[12,13]^ alveolar-capillary interface,^[14,15]^ and glomerular barrier.^[16]^ Studies to assess barrier structure and function are well-established and include fluorescence imaging of junctional proteins, molecular permeability measurements, and endpoint cell transmigration studies.^[17–21]^ Although open-well devices are convenient and feature tried-and-true experimental protocols, they cannot introduce fluid flows to mimic the perfused in vivo environment that circulates immune cells and contributes to barrier maturation.^[8,11,22]^ To address the lack of flow capabilities in conventional open-well assays, microfluidic platforms (e.g., microphysiological systems) that incorporate a porous culture membrane between the individually addressable top (luminal) and bottom (abluminal) fluidic channels have emerged.^[23–41]^

While microfluidic platforms offer distinct fluidic routing capabilities and feature dynamic control over biophysical and biochemical stimuli, they have not been widely adopted by bioscience laboratories where static, open-well assays remain the gold standard.^[10,11,42]^ This issue can partly be attributed to incompatibilities between microfluidic and conventional experimental workflows. For example, initial cell seeding in microfluidic channels is carried out under flow conditions; cells travel along the fluidic path (e.g., the tubing and channel) and finally settle onto the culture surface.^[11]^ Given the importance of the initial seeding density on barrier formation,^[43]^ this stochastic seeding approach poses a challenge because precisely controlling the uniformity of cells on the membrane under flow can be difficult to replicate between experiments. Conversely, open-well seeding simply adds a known cell number directly to the culture membrane to achieve the desired density (i.e., cells.cm^-2^). Although many standard assays can now be performed in a microfluidic channel under controlled flow conditions (e.g., immunostaining, protein isolation, molecular permeability, and RNA extraction), the sample handling and experimental protocols are naturally more complex and less familiar than those used for static, open-well formats, and require significant optimization efforts.^[2,11]^ An ideal system would unite the distinct advantages of open-well and microfluidic techniques and provide an opportunity to switch the configuration during an experiment based on the experimental requirements and user preference.

Here, we introduce a reconfigurable platform that combines the ease-of-use and standard protocols found in open-well formats with the controlled dynamic flow capabilities provided by microfluidics. Building from our open-well modular-μSiM (m-μSiM; **micro**physiological system enabled by a **si**licon **m**embrane) platform described in a companion paper,^[44]^ we use a tool-free magnetic latching approach to insert a microfluidic flow module onto an established endothelial barrier in the m-μSiM and introduce luminal flow for shear conditioning and leukocyte introduction. The m-μSiM features an ultrathin culture membrane with molecular level thinness (100 nm), high porosity (15%), and excellent live imaging capability that addresses problems with track-etched membranes materials used in commercial membrane-based culture systems (**Figure 1B**).^[26,29–31]^ As shown in our companion paper, barriers can be established in the m-μSiM using familiar open-well protocols with excellent reproducibility across engineering and bioscience-focused laboratories.^[44]^

To highlight the flexible nature of our approach, we seed and establish an endothelial monolayer in the open-well format and reconfigure the platform into a fluidic system. Using particle image velocimetry (PIV), we experimentally validate the computational model of the flow path and demonstrate the shear-induced alignment of endothelial cells. We then introduce leukocytes to the endothelial barrier under flow conditions that limit unwanted leukocyte shear-activation and visualize the different phases of cell recruitment, including rolling, arrest, probing, and transmigration across the endothelial monolayer in response to abluminal N-Formylmethionine-leucyl-phenylalanine (fMLP) stimulation. By enabling the facile reconfiguration of the open-well m-μSiM into the fluidic mode, we aim to provide bioscience and engineering-focused researchers a simple, modular workflow that provides on-demand flow capabilities.

## 2. Experimental Section

### Device Fabrication

#### Open-well m-μSiM device

100 nm ultrathin nanoporous silicon nitride membranes with 15% porosity and 60 nm pore sizes were provided by SiMPore Inc. (Rochester, USA). The overall dimensions of the membrane chip, including the silicon support, were 5.4 × 5.4 × 0.3 mm with a central permeable window of 2 mm × 0.7 mm × 0.1 μm. m-μSiM components, including the transparent COP (cyclic olefin polymer) base, PSA (pressure-sensitive adhesive) channel and support layers, and acrylic layers, were purchased from Aline Inc. (Signal Hill, USA). The membrane and m-μSiM components were assembled in a layer-by-layer manner, as described in a companion paper **(Figure 2A)**.^[44]^ The porous region of the membrane (yellow) is surrounded by a silicon support (blue) (**Figure 2B**). As shown in **Figure 2C**, the membrane separates the two compartments of the m-μSiM into luminal and abluminal sides while allowing soluble factor communication between compartments to occur over length scales consistent with the ∼100 nm thick basement membrane found in vivo.^[29–31]^

**Figure 2.**
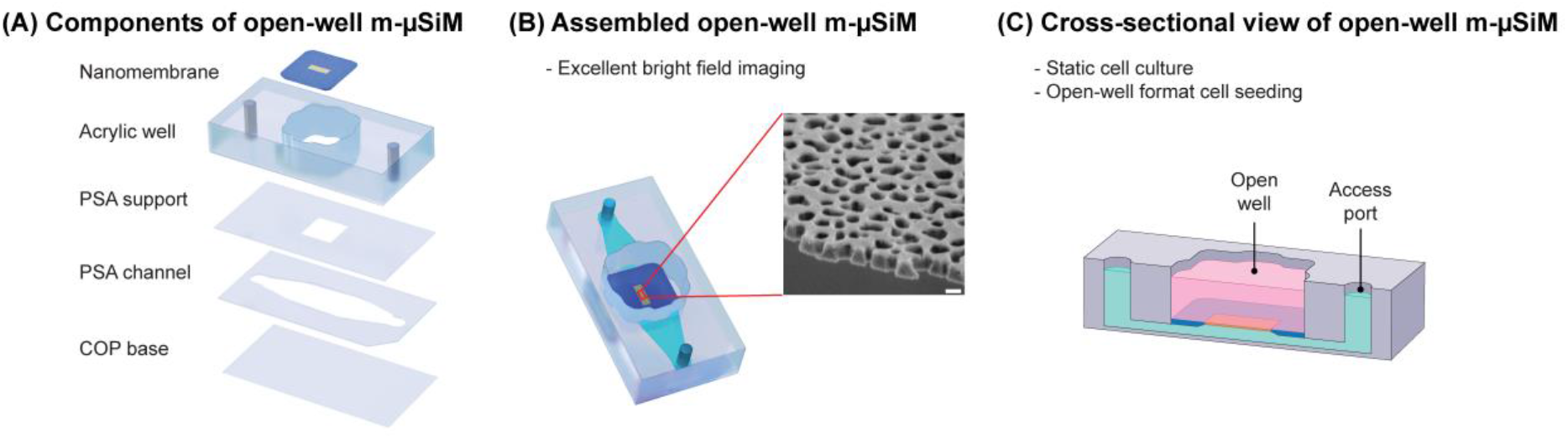
Schematic illustration of open-well m-μSiM. (A) Open-well m-μSiM consists of an acrylic top well and PSA microchannel separated by an ultrathin membrane. (B) Assembled view of the open-well m-μSiM showing the porous membrane (yellow) and silicon support (blue). The inset shows an SEM image of the 100 nm thick membrane. Scale bar = 100 nm. (C) A cross-sectional view of the open-well m-μSiM showing luminal (pink) and abluminal (green) compartments separated by the porous membrane.

#### Flow module

A polydimethylsiloxane (PDMS) flow module (molded microchannel dimensions, w = 1.5 mm, h = 0.2 mm, l = 5 mm) was fabricated using standard soft lithography techniques.^[45–47]^ Briefly, SU-8 3050 photoresist (Kayaku Advanced Materials, USA) was spin-coated onto a 100 mm diameter silicon wafer (University Wafers, USA), followed by a soft bake at 95 °C for 45 min. The photoresist was then exposed to UV light (250 mJ.cm^-2^) through a high-resolution printed photomask, baked for 5 min at 95 °C, and then developed in SU-8 developer until the features were resolved. The wafer was rinsed with IPA and dried using a stream of air. To fabricate flow modules with a clover shape and desired overall thickness (3.3 mm), an acrylic divider with a thickness of 3.3 mm was laser cut and attached to the wafer with PSA (MP467, 3M, USA) (**Figure S1A**). Degassed PDMS prepolymer (10:1 base to catalyst ratio) (Sylgard 184, USA) was poured over the mold and a ruler was used to level the upper surface before curing for 1 h at 70 °C on a hotplate. The flow modules were extracted from the mold and access ports were cored using a 1 mm biopsy punch (World Precision Instruments, USA).

#### Housings

Upper and lower housings were fabricated by laser-cutting acrylic sheets with a thickness of 2 and 3.5 mm, respectively. The lower housing was bonded to a coverslip using PSA. Nickel-plated magnets with a diameter of 4.75 mm and pull force of 0.34 Kg (K&J Magnetics, USA) were press fit into the housings.^[48,49]^

#### Patterning stencil

To create stencils with cone-shaped wells (h = 3.3 mm, taper angle = 57º), a custom aluminum mold was milled using a CNC machine and then polished. An acrylic divider with a thickness of 3.3 mm was laser cut and attached to the mold with PSA to obtain stencils with a clover-shaped footprint (**Figure S1B**). Degassed PDMS prepolymer (10:1 base to catalyst ratio) was poured over the mold and cured for 1 h at 70 °C on a hotplate.

#### Flow circuit

A custom-made flow circuit (**Figure S2**) was designed and fabricated using sample collection vials, Gauge 21 NT dispensing tips (Jensen Global, USA), 0.22 μm PVDF syringe Filters (Perkin Elmer, USA), and 0.5×0.86 Micro Flow tubes (Langer Instruments, USA). A peristaltic pump (Langer Instruments, USA) was used for flow circulation.

### Simulation

A steady-state 3D simulation of the fluid domain was performed using laminar flow physics in COMSOL Multiphysics®. Constant inlet velocity corresponding to a range of flow rates (Q=10, 100, 200, 500, and 1000 μl.min^-1^) was applied to the inlet, while the outlet boundary condition of pressure P=0 was applied to the outlet. Wall boundary conditions were applied to all other surfaces. The physics-controlled mesh was used and a mesh independence study was conducted. To improve the accuracy of the solution, a second-order discretization was utilized.

### Particle Image Velocimetry (PIV)

To validate the COMSOL model, experimental fluid flow analysis was carried out using particle image velocimetry on a Nikon Eclipse TE2000-S microscope equipped with a pulsed laser (TSI, MN, USA) (**Figure S3**). For this purpose, 5 μm fluorescent polystyrene latex particles (Magsphere, CA, USA) in deionized (DI) water were injected at defined flow rates (Q=10, 100, 200, 500, and 1000 μL.min^-1^) into the chip using a syringe pump. The PIV measurement was carried out on three devices and was repeated three times at each flow rate. All measurements were conducted at the middle of the channels (z=100 μm). For each measurement, 50 pairs of images were captured. The time interval between image pairs was varied from 100 to 1000 μs depending on the flow rate. The velocity was determined for each image pair using cross-correlation algorithms, and the results of the 50 instantaneous measurements were used to calculate the mean velocity for each replicate device. All data capture and analysis were performed using TSI Insight 4G™ software (TSI, MN, USA).

### Cell Culture

Human Umbilical Vein Endothelial Cells (HUVEC) (Thermo Fisher Scientific, USA) were cultured in EBM™-2 Basal Medium (Lonza, USA) supplemented with EGM™-2 Endothelial Cell Growth Medium-2 BulletKit™ (Lonza, USA). HUVECs were used between passages 3– 6. Cells were cultured in T25 flasks and were enzymatically dissociated using TrypLE (Thermo Fisher Scientific, USA) for 3 min and centrifuged at 150 G for 5 min, and re-suspended. For cell seeding, m-μSiM membrane was coated with 5 μg.cm^-2^ Fibronectin (Corning, USA) for 1 hour at room temperature and rinsed with cell media. A patterning stencil was inserted in the m-μSiM well and cell media was added to the well and bottom channel of the device. Next, cells were seeded at a density of 40,000 cells.cm^-2^ into the well and incubated for 24 hours at 37° C and 5% CO2. For the conventional open-well device, ThinCert™ transparent polyethylene terephthalate inserts (Greiner Bio-One, USA) were purchased, and a similar cell seeding procedure was carried out. To introduce shear flow to the cells, the m-μSiM was reconfigured into a fluidic system and cells were subjected to 10.7 dynes.cm^-2^ at a flow rate of 580 μL.min^-1^ for an additional 24 hours. Before use, the flow module, housings, and flow circuit were sterilized via exposure to UV light for 30 min. Next, PBS 1X (VWR, USA) was run through the flow circuit for 2 min. Then, reservoirs and tubes were filled with cell media, and the assembled device was connected to the flow circuit using Gauge 21 dispensing tips (Jensen Global, USA).

### Neutrophil Isolation and Transmigration

Polymorphonuclear neutrophils were isolated using a protocol described in Salminen et al., which was approved by the Research Subjects Review Board at the University of Rochester.^[29,51]^ Briefly, venous whole blood from consenting healthy donors was collected into sodium heparin-coated vacuum tubes and cooled to room temperature. Blood samples were carefully layered on top of equal volumes of 1-Step Polymorph solution (Accurate Chemical & Scientific Co., Westbury, NY) and centrifuged at 500g for 30 minutes. Post centrifugation, all density separation components (except for neutrophil rich layers) were discarded. Neutrophils were extracted into 15 ml conical tubes and washed twice in 10 mL of wash buffer (calcium and magnesium-free Hank’s balanced salt solution, 10 mM HEPES Sodium Salt, and 5 mg.mL^-1^ bovine serum albumin) via pelleting at 350g for 10 minutes, supernatant extraction, and subsequent resuspension. The washed neutrophil rich solution was then depleted of red blood cells via RBC lysis (4.5 mL of 1/6^th^ x PBS for one minute followed by 1.5 mL of 4x PBS for balancing) before one final wash with 10 mL of wash buffer. After the final wash, the neutrophil -rich pellet was resuspended in 1 mL of wash buffer, transferred to a 1.5 mL conical tube, and left in a rotating stand to prevent neutrophils from settling.

Neutrophils were introduced to the system for transmigration studies once a fully confluent monolayer of endothelial cells was formed. Isolated neutrophils from the resuspended stock were aliquoted into MCDB-131 complete medium (VEC Technologies, Rensselaer, NY) at 3 million cells.mL^-1^ and introduced into the top channel following a flow-stepping process of 10, 1, 0, 1, 10 μL.min^-1^ (30 seconds each until 10 μL.min^-1^ is reached again and maintained) using a syringe pump. 10 nM fMLP (N-Formylmethionyl-leucyl-phenylalanine) (Sigma Aldrich, St. Louis, MO) was incorporated into MCDB-131 media and introduced to the bottom channel of the device to serve as a neutrophil chemoattractant. Phase-contrast microscopy (Nikon Eclipse Ti2, Tokyo, Japan) at 40x was utilized to record a video (Andor Zyla sCMOS, Belfast, United Kingdom) at 20 Hz of rolling, arrest, probing, and transmigration events in the m-μSiM device. All neutrophil experiments were performed within 5 hours of blood draw.

### Flow Cytometry

Neutrophils were aliquoted into negative control, positive control, and flow module groups before antibody staining and flow cytometry. All aliquots consisted of 1 mL of neutrophil rich fluid and were deposited into 1.5 mL conical tubes. Negative control neutrophils were suspended in wash buffer at a concentration of 900k cells.mL^-1^. Positive control neutrophils were incorporated into MCDB-131 Complete Media at a concentration of 900k cells.mL^-1^ and supplemented with 10 ng.mL^-1^ IL-8 (rh CXCL8/IL-8, R&D Systems, Minneapolis, MN) for 15 minutes before rinsing and stained. Both control groups were left on a rotating stand to prevent neutrophils from settling. For the flow module group, neutrophil rich fluid from a non-agitated reservoir containing 3 million cells.mL^-1^ (suspended in MCDB-131 Complete) was pulled through the m-μSiM device at a flow rate of 10 μL.min^-1^ using a syringe pump (Chemyx Fusion 200, Stafford, TX) over 40 minutes. To remove potential effects of syringe geometry and associated shear stresses on neutrophil activation, 152 mm of 0.8 mm inner diameter silicone tubing (Model 95802-01, Cole Parmer, Vernon Hills, IL), placed between the syringe pump and flow module. This length held a volume of ∼300 μL neutrophil-rich media, which resulted in 900k cells after accounting for ‘loss’ from the volume of the fluidic circuit/device, and media from this section of tubing was exclusively used for analyzing flow module neutrophil activation. The samples were collected in separate 1.5 mL conical tubes, pelleted, and resuspended in 400 μL buffer (Flow Cytometry 1x Staining Buffer, R&D Systems, Minneapolis, MN). The 400 μL suspension for each experimental condition was then distributed into four 100 μL aliquots in 1.5 mL conical tubes representing the unstained control, isotype control, and L-selectin groups. Staining was performed according to manufacturer protocol using antibodies conjugated to Alexa Fluor 488 (R&D Systems, Minneapolis, MN). Flow cytometry was performed post live staining on a Guava EasyCyte benchtop machine. Sampling was set at 3000 gated events and data gathered was analyzed in FCS Express, GraphPad PRISM, and Wolfram Mathematica.

To assess neutrophil activation rates in the flow module, a histogram distance measurement (Jaccard Index) was calculated utilizing green fluorescence intensity data.^[52]^ As L-selectin is cleared from an activated neutrophil membrane surface via ectodomain shedding, decreased expression of L-selectin in an activated neutrophil population will result in greater rates of non-specific binding for anti-L-selectin staining antibodies.^[53–55]^ Activated neutrophil populations displayed higher degrees of overlap between respective L-selectin expression histograms and isotype control histograms. Higher degrees of neutrophil activation corresponded to higher Jaccard Index values, which were calculated for each experimental group.

### Immunostaining and Imaging

For actin and nuclei staining, cells were fixed in 4% paraformaldehyde (Fisher Scientific, USA) for 15 min. Then, cells were permeabilized in Triton X-100 (0.1%) for 10 min and washed with PBS Tween-20 (PBST). Next, cells were blocked in 40 mg.mL^-1^ BSA (Alfa Aesar, USA) for 30 min at room temperature. Cells were labeled with Hoechst 33342 (300 nM) (Molecular Probes, USA) for 10 min and AlexaFluor 488 conjugated phalloidin (ThermoFisher Scientific, USA) (1:400) for 15 min to visualize nuclei and actin fibers, respectively. Finally, cells were washed with PBS Tween-20 and stored in PBS. For viability assays, the LIVE/DEAD™ Cell Imaging Kit (Thermo Fisher Scientific, USA) was used based on the vendor’s protocol. In the case of imaging cells on the silicon support of the membrane, a coverslip was placed on the m-μSiM and the device was flipped and imaged from top to bottom. Phase and fluorescence imaging of cell samples was performed on an Olympus IX-81 inverted microscope with CellSens software (Olympus, Japan). All image collection settings were consistent across experimental sets to allow comparison.

### Customization of m-μSiM and introduction of aligned ECM fibers

A customized “segmented” channel geometry (wide segment: L = 5mm, W = 4mm, narrow segment: L = 5 mm, W = 1 mm) was fabricated by laser-cutting a 130 μm PSA sheet. During the assembly of the m-μSiM, the standard bottom channel was replaced with the customized channel. Atelo-bovine collagen type-1 (Nutragen, Advanced BioMatrix, CA, USA) with a final concentration of 2.5 mg.mL^-1^ was prepared as described previously.^[46,50]^ The diluted solution was injected into the custom-designed bottom channel of the m-μSiM through a Gauge 20 IT dispensing tip at a flow rate of 200 μl.min^-1^ using a syringe pump. The device was maintained at 37°C in an incubator for one hour to induce collagen polymerization. Collagen fibers were imaged using a Leica SP5 confocal microscope (Leica Microsystems, Germany) with a 40X water objective (HCX PL APO 40x/1.1 W) at 1.7x optical zoom. At each location, stacks with a thickness of 2 μm comprising 13 slices were captured. Then, stacks were merged and projected using a maximum projection FIJI (NIH, USA). LOCI CT-FIRE was used to identify collagen fibers in the confocal reflectance microscopy (CRM) image and calculate the coefficient of alignment (CoA).^[46,50,56]^

### Image analysis

Image analysis was carried out using ImageJ (National Institutes of Health, USA). For cell alignment quantification, the Analyze Particles module in Image J was used and results were plotted as radar graphs using MATLAB CircHist plugin. ^[57]^

## 3. Results

### 3.1. Functional module provides microfluidic capabilities to static, open-well m-μSiM

In our companion paper,^[44]^ we demonstrated that the open-well m-μSiM platform could support the formation and maturation of a blood-brain barrier mimetic using static cultures of iPSC-derived endothelial cells in different laboratory environments (engineering and bioscience laboratories) using mass-produced components and a common culture protocol. To expand the experimental capabilities of the m-μSiM platform, we demonstrate that the open-well format can be reconfigured into a fluidic device using magnetic assembly.^[48–50]^ As shown in **Figure 3A**, the core m-μSiM is first placed into a cavity defined by a lower housing and a coverslip. A molded “flow module” containing a lithographically defined channel and access ports (**Figure S4**) is added to the open-well reservoir. The flow module is sealed against the silicon support layer using magnetic “latching” produced by the attraction force from magnets embedded in the top and bottom housing layers (**Figures 3B, C, <u>Video S1</u>**). The matching “clover” shaped geometry of the flow module and reservoir ensured that the microchannel features were positioned without damaging the membrane.

**Figure 3.**
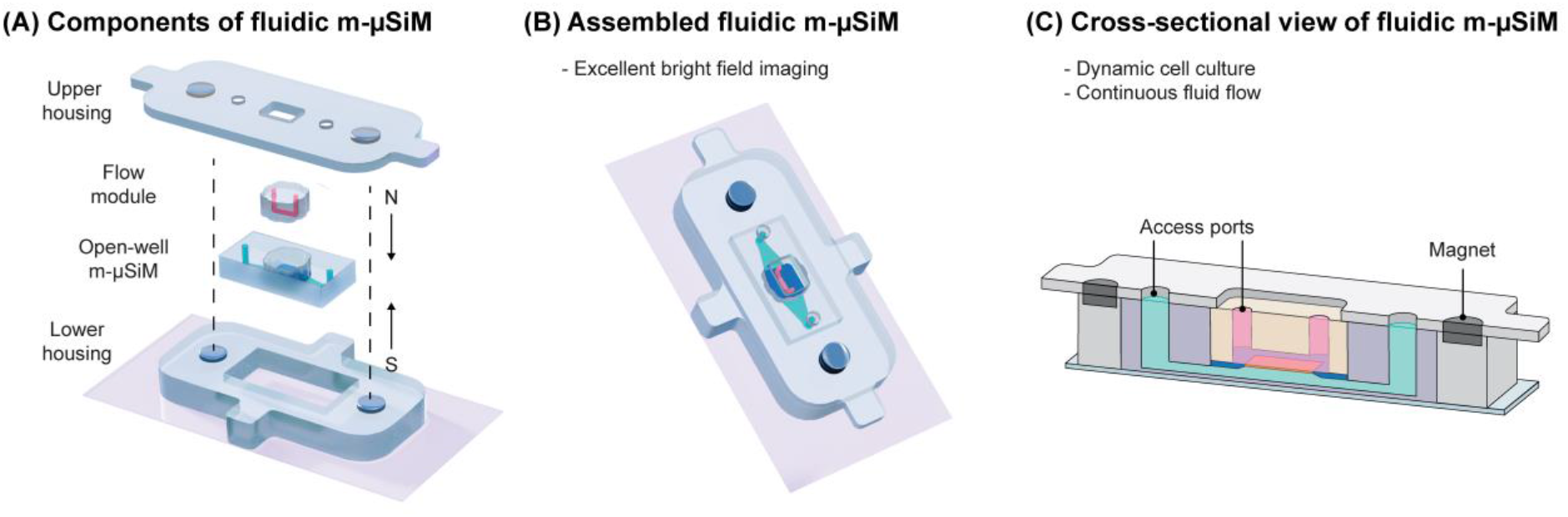
Schematic illustration of modular fluidic m-μSiM. (A) The open-well m-μSiM is reconfigured into the fluidic m-μSiM using magnetic sealing of a flow module by upper and lower housings with embedded magnets. (B) Assembled view of the fluidic m-μSiM showing fluid in the flow module channel above (pink) and below (green) the membrane. (C) Cross-sectional view of the reconfigured fluidic m-μSiM with the flow path over the membrane identified.

Although it is not a typical consideration for conventional static open-well platforms, the cell seeding approach must be carefully considered when developing a platform designed to be reconfigured between open-well and fluidic modes. For example, as shown in **Figure 4A**, during cell seeding into the open-well m-μSiM, the HUVECs settled onto the membrane and surrounding silicon support and formed a continuous monolayer. When the flow module was magnetically sealed, cells outside of the microchannel boundary compressed and appeared non-viable (red), while cells inside the channel remained viable (green). Damage to the monolayer around the periphery of the channel may be problematic because the overall integrity of the barrier could be compromised and confound experimental findings. To alleviate such concerns, we designed a removable stencil that fits within the m-μSiM well and defines a patterning region where cells settle preferentially onto the membrane surface. Although there is an additional step in the workflow, cells are localized to a region within the microchannel boundary and cell damage does not occur after reconfiguration (**Figure 4B**).

**Figure 4.**
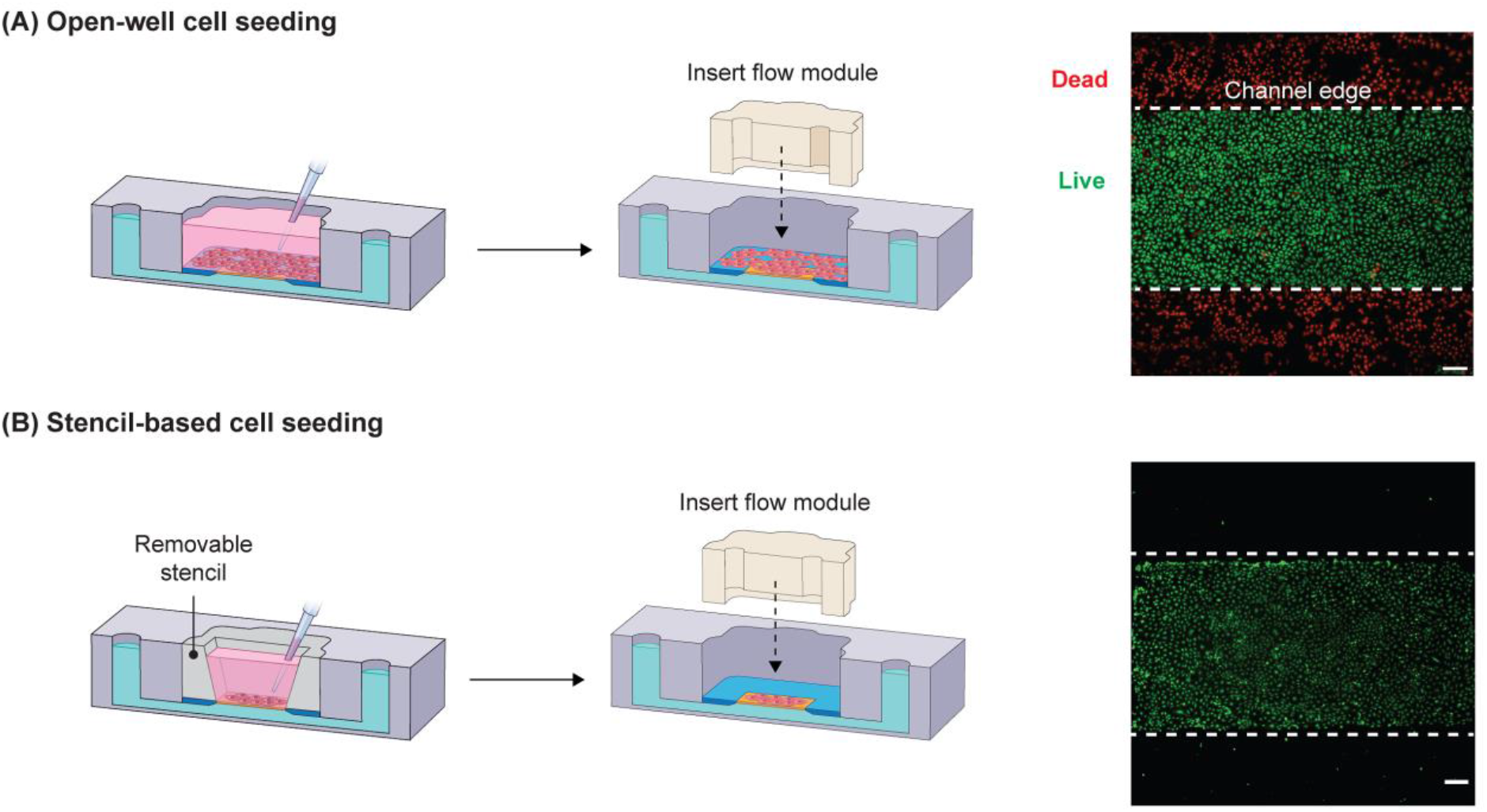
Comparison of open-well cell seeding vs stencil-based cell seeding. (A) Open-well format cell seeding results in cell settlement all over the well surface and consequent establishment of a monolayer on the membrane as well as its surrounding silicon chip. In this scenario, cells on the membrane’s surroundings are damaged upon inserting the flow module. (B) Whereas, in stencil-based seeding, cells are positioned selectively; hence, the monolayer only forms on the membrane surface and there are no cells in the surrounding of the membrane to be damaged upon inserting the flow module. Scale bars = 200 μm.

The placement of the stencil within the m-μSiM is compatible with conventional cell seeding protocols and is the first step in the experimental workflow **(Figure 5A, B)**. After patterning and formation of a confluent cell monolayer, the user has two options: (i) for static cultures, the device can be used as-is in the open-well platform (**Figure 5C**), (ii) for flow experiments, the device can be reconfigured into the fluidic mode as shown in **Figure 5D**. The reversible magnetic latching mechanism supports on-demand reconfiguration between the open-well and fluidic formats (**<u>Video S1</u>**). For example, after flow stimulation, the device can be reconfigured to the open-well format and the user has the option to perform assays including immunostaining and barrier permeability measurements using standard experimental protocols as described in the companion paper.^[44]^ The user also has the option to keep the channel in place and use microfluidic assay protocols if desired.

**Figure 5.**
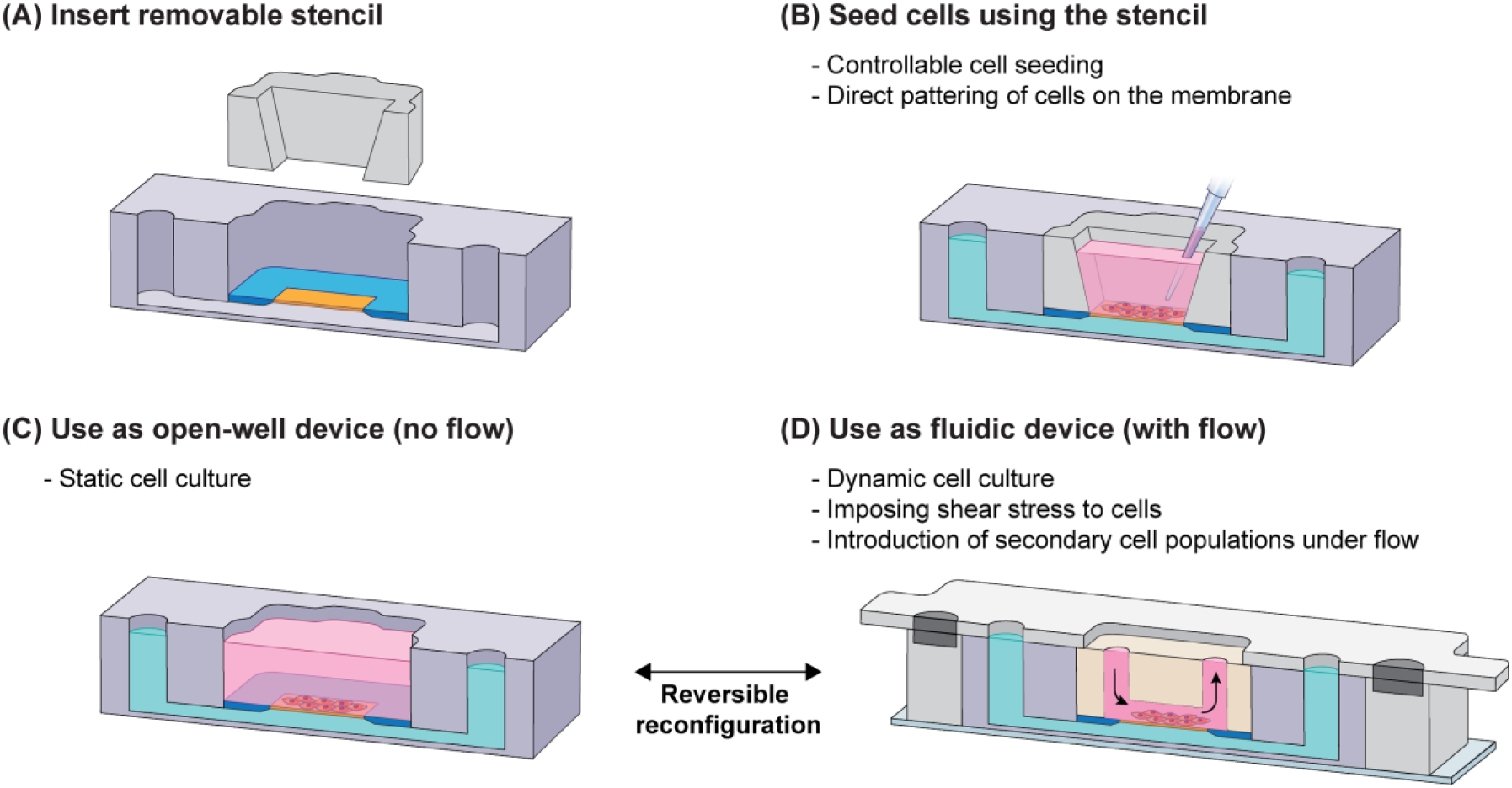
Experimental workflow for the fluidic m-μSiM. (A) Before cell seeding, a removable stencil is inserted into the m-μSiM well; then, (B) cells can be directly patterned on the membrane to prevent the distribution of cells on the membrane surrounding and consequent cell damage upon insertion of the flow module. (C) After the monolayer is established, the stencil can be kept or removed and the device can be used for static cell culture in the open-well format. (D) If desired, the user can reconfigure the device into a fluidic mode by inserting the flow module in the m-μSiM well and sealing it magnetically. After the flow experiment, the housings and flow module can be removed to reconfigure the system to the open-well format with direct access to the cells. The pink and green domains represent fluid in the vessel side and the tissue side of the barrier, respectively.

### 3.2. Computational modeling and experimental validation of flow characteristics in the fluidic m-μSiM

Flow-induced shear stress is a well know biophysical cue that positively influences the structure and function of vascular barriers (e.g., endothelial alignment, barrier integrity, and permeability).^[8,58]^ To characterize the flow within the device, we developed a COMSOL model and experimentally validated the simulation using particle image velocimetry (PIV). Our results showed that the simulated and measured velocities matched within 10% (n=3), with excellent agreement over the tested range (linear regression, R^2^=0.99) **(Figure 6A,B)**. Since there was a match between the experimental and simulated velocities, COMSOL results were used as a guide to calculate fluid-induced shear stresses at the cell monolayer and identify maximum shear stress values along the flow path (**Table S1**).

**Figure 6.**
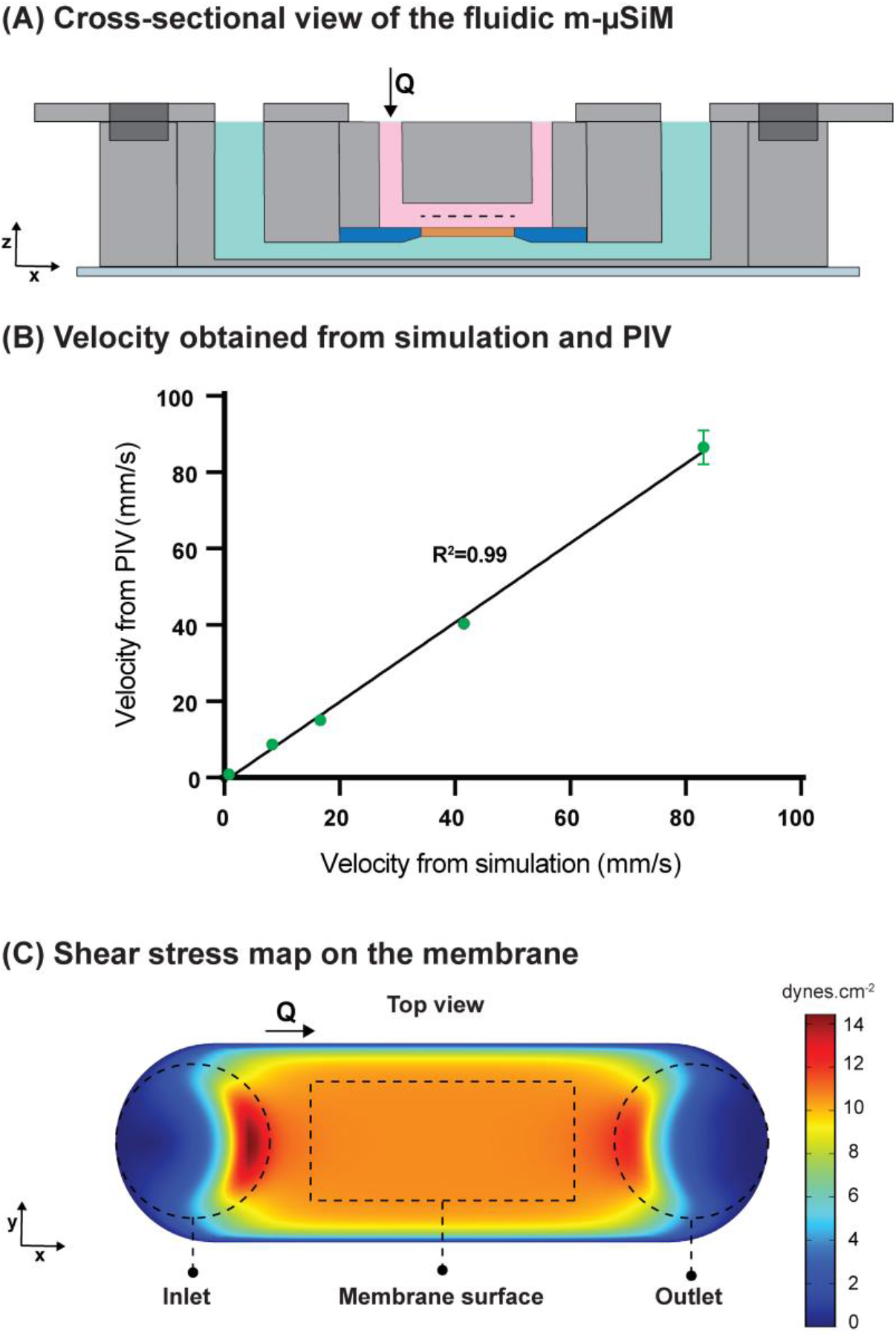
(A) Schematic of the m-μSiM with attached flow module. The flow path is shown in pink. The velocities from the simulation and PIV were obtained from the mid-plane of the microchannel at z = 100 μm (dashed line) at Q = 10, 100, 200, 500, 1000 μL.min^-1^. (B) The comparison of velocities from the simulation and PIV measurements (R^2^ = 0.99, n = 3) shows excellent agreement. (C) The shear stress map in the flow channel confirms that the flow channel geometry results in a uniform shear stress distribution at the membrane surface (denoted by the dashed rectangular region).

Endothelial cells are exposed to fluid-induced shear stresses in vivo and their elongation along the flow direction results in enhanced barrier function.^[8,30]^ With ∼10 dynes.cm^-2^ recognized as the typical threshold required for the alignment of endothelial cells,^[30,59,60]^ we performed a parametric study using COMSOL to determine the flow rate required to produce a wall shear stress of ∼10 dynes.cm^-2^ in our flow module. Based on the simulation results, we selected 580 μL.min^-1^ as the inlet flow rate, corresponding to 10.7 dynes.cm^-2^ shear stress at the membrane surface. We mapped the shear stress distribution at the endothelial surface and confirmed that there we no spatial variations along the monolayer as shown in **Figure 6C**.

To confirm that the predicted flow rate was sufficient to align cells, the open-well m-μSiM was reconfigured to the fluidic m-μSiM after initial monolayer formation, and cells were exposed to 10.7 dynes.cm^-2^ of shear stress (i.e., Q = 580 μL min^-1^). Consistent with the literature,^[30,59,60]^ shear-stimulation resulted in cell alignment (89±4.9 %) along the flow direction **(Figure 7A)**, while cells in the static culture control (no flow) remained randomly orientated (**Figure 7B**).

**Figure 7.**
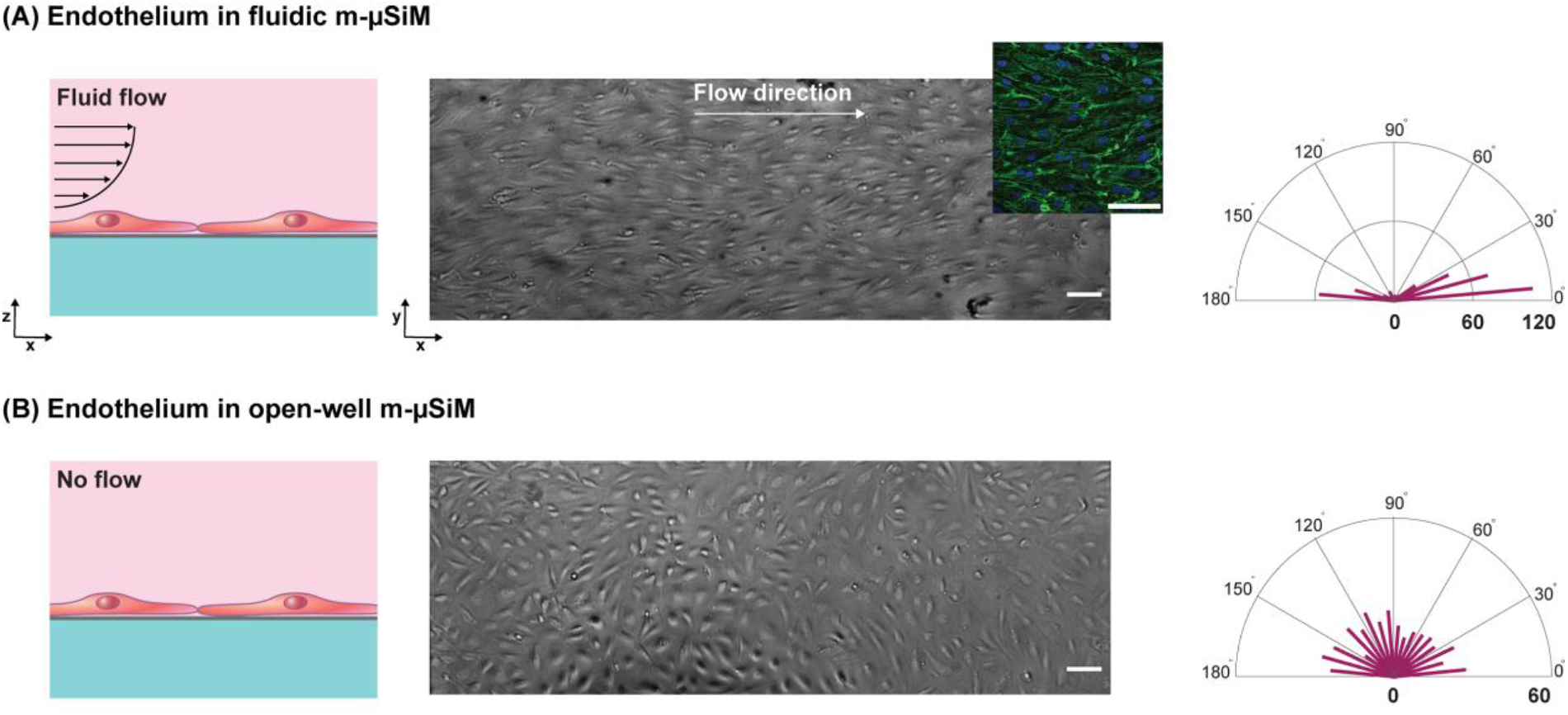
Comparison of endothelial cells cultured in open-well and fluidic configurations of the m-μSiM. First, cells were seeded and maintained in open-well format for 24 hours to establish a confluent monolayer in m-μSiM devices. Then, one of the devices was reconfigured into the fluidic mode to induce shear alignment. (A) Cells cultured under dynamic conditions showed alignment along the flow direction due to continuous exposure to 10.7 dynes.cm^-2^ shear stress at a flow rate of 580 μL.min^-1^. The inset shows actin and nuclei of aligned cells in green and blue, respectively. (B) Cells cultured under static culture showed no alignment and were randomly oriented in different directions due to lack of shear stress. Radar plots show quantification of cell alignment with respect to the flow direction (x-axis). The length of each bar represents the number of cells in the corresponding direction. Scale bars = 100 μm.

During tissue injury or inflammation, leukocytes (e.g., neutrophils) are recruited from the bloodstream to the endothelial surface in response to inflammatory factors and then transmigrate across the barrier in a coordinated process.^[29,31,58]^ It is important to note that neutrophils can become activated upon exposure to shear stress greater than ∼1.5 dynes.cm^-2^.^[31,61,62]^ Shear activation can change neutrophil responsiveness to inflammatory factors and reduce transmigration efficiency.^[55]^ Thus, to limit shear activation effects within the flow module, we mapped the shear stress distribution along the flow path using our PIV-validated COMSOL model and selected a flow rate where the maximum shear stress was less than the activation threshold of 1.5 dynes.cm^-2^. For the neutrophil introduction, Q = 10 μL.min^-1^ was selected to both minimize shear exposure and ensure adequate deposition onto the membrane surface at the initial stage of the transmigration process (**Figure S5**). **Figure 8A** shows that the maximum predicted shear stress in the flow path (0.4 dynes.cm^-2^) occurs at the intersection of vertical access ports and the horizontal channel and is below the shear activation threshold.

**Figure 8.**
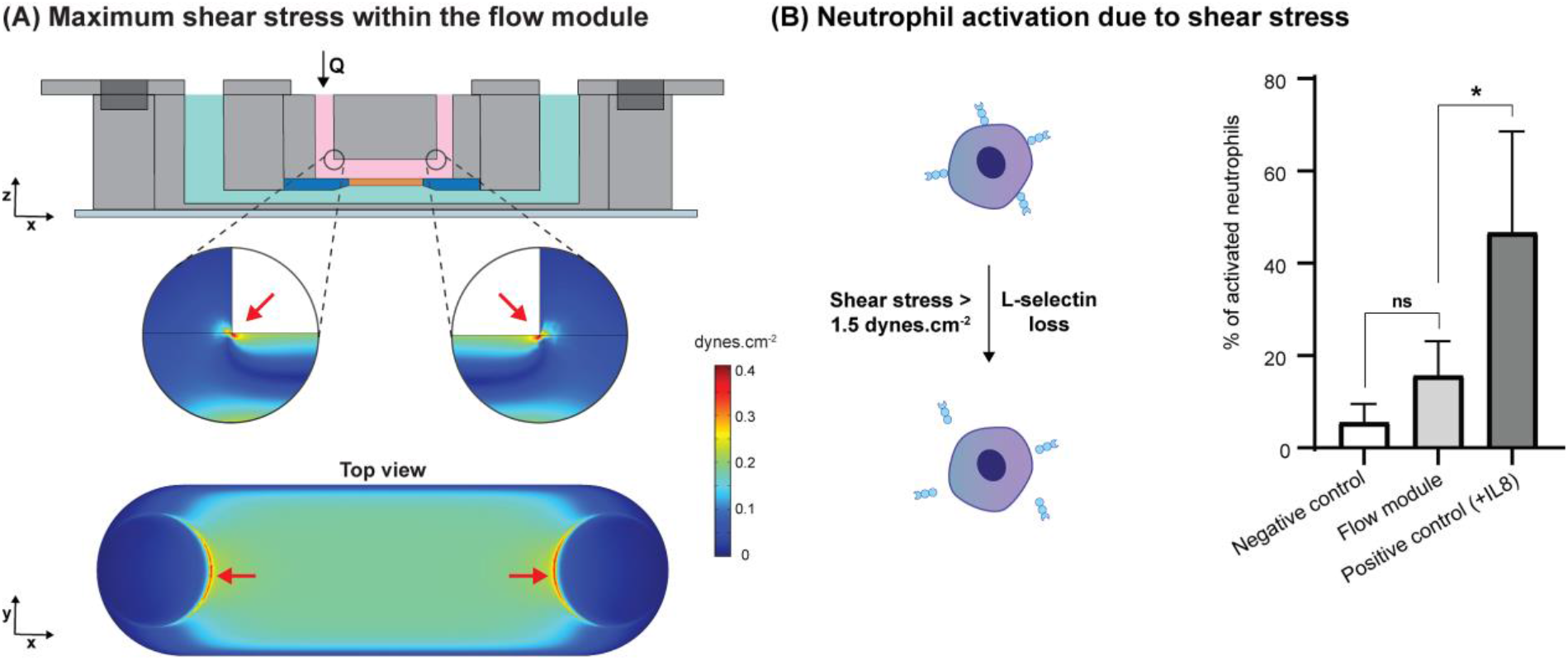
Shear-induced activation of neutrophils. (A) Maximum shear stress within the flow module occurs at the intersection of vertical access ports and horizontal microchannel (red arrows); based on the simulation, the maximum shear stress is 0.4 dynes.cm^-2^ at an inlet flow rate of 10 μL.min^-1^ and neutrophils should not become activated by the fluid dynamics within the flow module. (B) Upon exposure of neutrophils to shear stresses above 1.5 dynes.cm^-2^, they become activated and shed the surface marker, L-selectin. Quantitative analysis of neutrophil activation based on L-selectin loss shows that the percentage of activated neutrophils in the negative control and flow module are not statistically different, while the positive control with neutrophils treated with 10 ng.ml^-1^ of the cytokine IL8 shows a significant statistical difference (p-value = 0.037, n = 4).

To confirm our model-based prediction that neutrophils introduced at Q=10 μL.min^-1^ should not exhibit shear activation in the flow module, we carried out a flow cytometry study in which the loss of the surface marker, L-selectin, was used as a metric to assess activation (**Figure 8B**).^[53–55]^ Our results showed no statistical differences between the percentage of activated neutrophils in the population collected from the flow module and the negative control group (**Figure 8B, Figure S6**), while significant statistical differences were observed compared to the IL8-treated positive control condition (p-value = 0.037, n = 4). Thus, we concluded that our validated model could be used to select flow rates to limit shear activation during the introduction of neutrophils under flow conditions.

### 3.3. Visualization of the neutrophil transmigration process under flow

Live monitoring of the neutrophil recruitment and transmigration process can help provide a deeper understanding of immune cell interactions with the vascular barrier in a physiological setting.^[29,31]^ Using live imaging and the fluid flow capabilities of our platform, we captured the distinct phases of neutrophil transmigration across the endothelial barrier in response to the chemotactic factor fMLP introduced to the abluminal channel (**<u>Video S2</u>**). Upon introduction, neutrophils roll on the cell monolayer **(Figure 9A)**, then become arrested and probe the monolayer before transmigrating across the barrier (**Figure 9B**,**C**). As the neutrophil begins transmigrating, its appearance changes from phase bright (**Figure 9D**) to phase dark (**Figure 9E**).^[29,51]^ Such studies could not be conducted without an optically transparent membrane and fluid flow capabilities. Thus, our platform provides an opportunity to investigate the dynamics of leukocyte trafficking under flow to understand their role in inflammation using a simple bright-field imaging approach that is not possible using conventional open-well methods.

**Figure 9.**
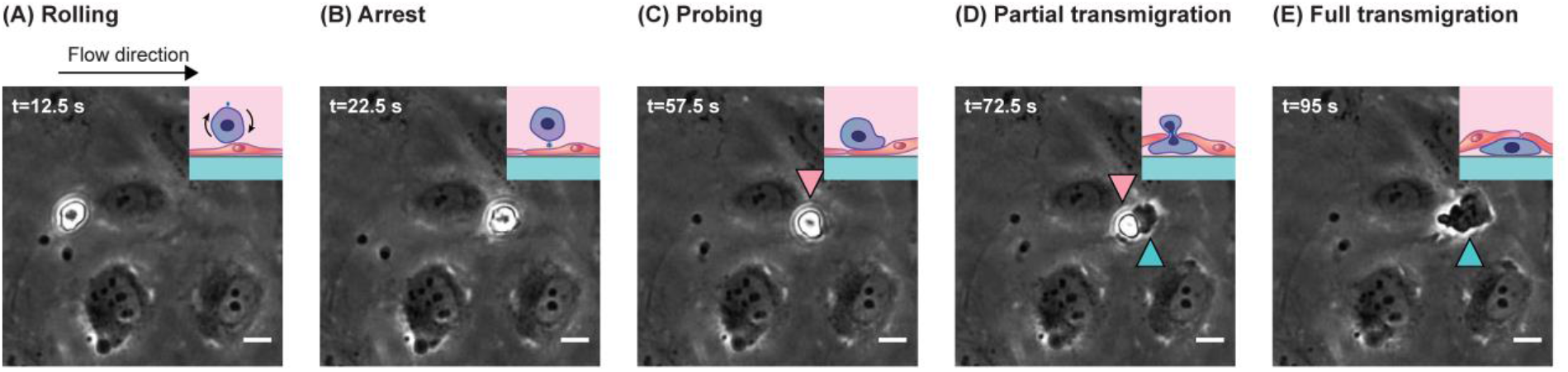
Bright-field imaging of neutrophil transmigration across an endothelial monolayer in response to abluminal fMLP stimulation. The HUVEC monolayer was established in the open-well format before the reconfiguration into the fluidic mode. Neutrophils were introduced under flow (left to right) and imaged to show the following steps: (A) rolling along the endothelial cell layer, (B) arrest, (C) probing their environment, and (D, E) transmigrating from the luminal side (pink arrowhead) to the abluminal side of the monolayer (green arrowhead). Scale bars = 10 μm.

### 3.4. Customization of the m-μSiM to enhance model complexity

The mass-produced components of the m-μSiM described in the companion paper have the potential to be customized at the individual laboratory level to enhance the physiological relevance of the barrier tissue model. For example, type-I collagen is of great interest as an extracellular matrix (ECM) material due to its unique microarchitecture and presence in various tissue environments.^[63]^ The incorporation of ECM gels into the abluminal side can enable users to study the post-transmigration guidance of leukocytes in the tissue environment. Our lab has previously shown that local extensional flows can induce long-range collagen fiber alignment.^[50]^ To obtain aligned collagen fibers below the membrane area, we customized the bottom channel of the m-μSiM to include a “segmented” channel geometry and replaced the standard component. As shown in **Figure S7**, we were able to obtain a 3D collagen gel with aligned fibers below the membrane in the m-μSiM. This simple demonstration highlights how components of m-μSiM can be easily redesigned to introduce new experimental capabilities in different laboratories.

## 4. Discussion

The goal of this work is to provide an experimental workflow to introduce flow capabilities to the open-well m-μSiM platform. Here, we use a magnetic latching approach that enables reconfiguration between open-well and fluidic modes during the experimental workflow to combine the unique advantages of each approach. In our platform, the barrier is established in the familiar open-well format and reconfigured to a fluidic system during an experiment by adding a functional flow module. This reconfiguration allows i) the monolayer to experience fluid-induced shear stress that models the physiological environment and ii) supports the introduction of leukocytes to the endothelial monolayer under flow conditions to visualize discrete steps in the recruitment process, including rolling, arrest, probing, and transmigration. The reconfiguration capability enables users to use established conventional protocols during experiments and also benefit from microfluidic capabilities when desired.

The reconfiguration capability requires a change in the workflow because the addition of the flow module can damage cells that are seeded uniformly over the membrane and silicon support. Here, we use a cell-seeding stencil that enables controlled positioning of cells directly onto the membrane surface that supports efficient seeding of cells at a known density, which is important because the starting cell concentration influences vascular barrier formation.^[43]^ The tapered well within the stencil increases media capacity (from 10 μl to 30 μl) while limiting bubble trapping that can occur within high aspect ratio, straight-walled reservoirs.^[64]^ Thus, cells can be maintained in long-term culture with the stencil in place if desired. The m-μSiM enables controlled cell seeding in the open-well format before the addition of the flow module and addresses challenges including cell losses along the flow path intrinsic to conventional microfluidic platforms and thus reduces the number of cells used.

Magnetic latching is a unique feature and provides a simple, tool-free assembly approach that allows on-demand switching between open-well and fluidic formats. As a result, different steps of an experiment, including permeability measurements and immunostaining can be conducted in the open-well format using established techniques that most laboratories are familiar with.^[44]^ To show the compatibility of our platform with multi-step immunostaining protocols, we removed the flow module after aligning HUVECs and stained for actin and DAPI in open-well format to visualize the cytoskeleton and nuclei, respectively (**Figure 6A**). Furthermore, we have obtained similar shear-induced alignment of HUVECs using the reconfigurable approach between our engineering laboratories (Rochester Institute of Technology and University of Rochester) using common components and showing excellent inter-laboratory reproducibility using the flow module. Although we used HUVECs in our experiments as a proof-of-concept, the m-μSiM is compatible with other cell populations including iPSC derived endothelial cells in co-cultured environments.^[44]^ Furthermore, the reversible latching mechanism offers the potential to add other functional modules including TEER measurement and passive pumping modules. We have also recently developed a miniaturized portable fluidic system featuring a miniaturized 3D-printed pressure regulator that can provide on-board flow control in future iterations of the platform.^[65]^

Our companion paper highlights reproducible results using the mass-produced m-μSiM, and here we show reconfiguration capabilities by adding a flow module and customization of the bottom channel of the m-μSiM. Although PDMS was used for the fabrication of the flow module, materials such as moldable polyurethane can be used if problems related to small-molecule partitioning are identified.^[34,66,67]^ Our components including, the flow module, stencil, and housings are also amenable to scaled up fabrication similar to the mass-produced components of the open-well m-μSiM.^[44]^ We anticipate that the reconfiguration capabilities presented here will allow more widespread use of flow-enhanced microfluidic barrier models in both bioscience and engineering laboratories.

## 5. Conclusion

To combine the advantages of open-well and microfluidic devices, we developed a functional flow module that enables on-demand reconfiguration of open-well m-μSiM into a fluidic mode. This reconfigurable platform provides not only conventional open-well cell seeding and barrier establishment but also microfluidic capabilities to introduce controlled flows and secondary cell populations. This unique platform utilizes reversible magnetic latching, allowing the flow module to be removed and the barrier directly accessed for downstream experiments. Users have the flexibility to easily switch between different device modes during an experiment and carry out each experimental step in their preferred mode. This platform includes open-well format cell seeding, fluid flow, live imaging, and direct access to the barrier, whereas all prior platforms lack one or more of these features. Since users can select the device format and protocols they are most comfortable with, we anticipate that this device will enable more widespread use of flow-enhanced barrier models in both engineering and bioscience laboratories.

## Supporting Information

Supporting Information is available in the supplement document.

## Acknowledgments

This research was funded by the National Institute of Health under award number 1R43GM137651, 1R61HL154249, and UG3TR003281. The authors thank Xian Boles from the RIT Medical Illustration MFA program for illustration support and RIT Machine Shop for aluminum mold fabrication.

## References

[1] D. Vera, M. García-Díaz, N. Torras, M. Álvarez, R. Villa, E. Martinez, ACS Appl Mater Interfaces 2021, 13, 13920.

[2] F. Yu, N. D. Selva Kumar, D. Choudhury, L. C. Foo, S. H. Ng, Drug Discovery Today 2018, 23, 815.

[3] L. Claesson-Welsh, E. Dejana, D. M. McDonald, Trends in Molecular Medicine 2021, 27, 314.

[4] P. Rajendran, T. Rengarajan, J. Thangavel, Y. Nishigaki, D. Sakthisekaran, G. Sethi, I. Nishigaki, Int J Biol Sci 2013, 9, 1057.

[5] M. Xu, G. Chen, W. Fu, M. Liao, J. A. Frank, K. A. Bower, S. Fang, Z. Zhang, X. Shi, J. Luo, Toxicological Sciences 2012, 127, 42.

[6] D. A. Chistiakov, A. N. Orekhov, Y. V. Bobryshev, Frontiers in Physiology 2015, 6.

[7] M. D. Sweeney, K. Kisler, A. Montagne, A. W. Toga, B. V. Zlokovic, Nat Neurosci 2018, 21, 1318.

[8] M. A. Kaisar, R. K. Sajja, S. Prasad, V. V. Abhyankar, T. Liles, L. Cucullo, Expert Opinion on Drug Discovery 2017, 12, 89.

[9] D. E. Ingber, Advanced Science 2020, 7, 2002030.

[10] Prashanth, H. Donaghy, S. P. Stoner, A. L. Hudson, H. R. Wheeler, C. I. Diakos, V. M. Howell, G. E. Grau, K. J. McKelvey, Cancers 2021, 13, 955.

[11] M. A. Kaisar, V. V. Abhyankar, L. Cucullo, in Blood-Brain Barrier (Ed: T. Barichello), Springer, New York, NY, 2019, pp. 55–70.

[12] N. L. Stone, T. J. England, S. E. O’Sullivan, Frontiers in Cellular Neuroscience 2019, 13.

[13] K. Hatherell, P.-O. Couraud, I. A. Romero, B. Weksler, G. J. Pilkington, Journal of Neuroscience Methods 2011, 199, 223.

[14] J. Kasper, M. I. Hermanns, C. Bantz, M. Maskos, R. Stauber, C. Pohl, R. E. Unger, J. C. Kirkpatrick, Part Fibre Toxicol 2011, 8, 6.

[15] H. Janga, L. Cassidy, F. Wang, D. Spengler, S. Oestern-Fitschen, M. F. Krause, A. Seekamp, A. Tholey, S. Fuchs, Journal of Cellular and Molecular Medicine 2018, 22, 982.

[16] T. Moriyama, K. Sasaki, K. Karasawa, K. Uchida, K. Nitta, Journal of Cellular Physiology 2017, 232, 3565.

[17] J. P. Gleeson, H. Q. Estrada, M. Yamashita, C. N. Svendsen, S. R. Targan, R. J. Barrett, International Journal of Molecular Sciences 2020, 21, 1438.

[18] T. J. Pell, M. B. Gray, S. J. Hopkins, R. Kasprowicz, J. D. Porter, T. Reeves, W. C. Rowan, K. Singh, K. B. Tvermosegaard, N. Yaqub, et al., SLAS DISCOVERY: Advancing the Science of Drug Discovery 2021, 26, 909.

[19] Merino, M. Sablik, S. S. Korevaar, C. López-Iglesias, M. Ortiz-Virumbrales, C. C. Baan, E. Lombardo, M. J. Hoogduijn, Frontiers in Immunology 2021, 12.

[20] E. S. Wittchen, R. A. Worthylake, P. Kelly, P. J. Casey, L. A. Quilliam, K. Burridge, Journal of Biological Chemistry 2005, 280, 11675.

[21] C. Wewer, A. Seibt, H. Wolburg, L. Greune, M. A. Schmidt, J. Berger, H.-J. Galla, U. Quitsch, C. Schwerk, H. Schroten, et al., J Neuroinflammation 2011, 8, 51.

[22] D. E. Ingber, Nat Rev Genet 2022, 1.

[23] F. R. Walter, S. Valkai, A. Kincses, A. Petneházi, T. Czeller, S. Veszelka, P. Ormos, M. Deli, A. Dér, Sensors and Actuators B: Chemical 2016, 222, 1209.

[24] Y. I. Wang, H. E. Abaci, M. L. Shuler, Biotechnology and Bioengineering 2017, 114, 184.

[25] J. D. Wang, E.-S. Khafagy, K. Khanafer, S. Takayama, M. E. H. ElSayed, Mol. Pharmaceutics 2016, 13, 895.

[26] R. Booth, H. Kim, Lab Chip 2012, 12, 1784.

[27] T. S. Frost, L. Jiang, R. M. Lynch, Y. Zohar, Micromachines 2019, 10, 533.

[28] S.-H. Chang, P.-L. Ko, W.-H. Liao, C.-C. Peng, Y.-C. Tung, Micromachines 2021, 12, 406.

[29] T. Salminen, J. Zhang, G. R. Madejski, T. S. Khire, R. E. Waugh, J. L. McGrath, T. R. Gaborski, Small 2019, 15, 1804111.

[30] T. S. Khire, A. T. Salminen, H. Swamy, K. S. Lucas, M. C. McCloskey, R. E. Ajalik, H. H. Chung, T. R. Gaborski, R. E. Waugh, A. J. Glading, et al., Cell Mol Bioeng 2020, 13, 125.

[31] Mossu, M. Rosito, T. Khire, H. Li Chung, H. Nishihara, I. Gruber, E. Luke, L. Dehouck, F. Sallusto, F. Gosselet, et al., J Cereb Blood Flow Metab 2019, 39, 395.

[32] K. J. Regehr, M. Domenech, J. T. Koepsel, K. C. Carver, S. J. Ellison-Zelski, W. L. Murphy, L. A. Schuler, E. T. Alarid, D. J. Beebe, Lab Chip 2009, 9, 2132.

[33] M. Zhou, X. Zhang, X. Wen, T. Wu, W. Wang, M. Yang, J. Wang, M. Fang, B. Lin, H. Lin, Sci Rep 2016, 6, 31771.

[34] M. Ishahak, J. Hill, Q. Amin, L. Wubker, A. Hernandez, A. Mitrofanova, A. Sloan, A. Fornoni, A. Agarwal, Frontiers in Bioengineering and Biotechnology 2020, 8.

[35] D. Huh, B. D. Matthews, A. Mammoto, M. Montoya-Zavala, H. Y. Hsin, D. E. Ingber, Science 2010, 328, 1662.

[36] P. Loskill, T. Sezhian, K. M. Tharp, F. T. Lee-Montiel, S. Jeeawoody, W. Mae Reese, P.-J. H. Zushin, A. Stahl, K. E. Healy, Lab on a Chip 2017, 17, 1645.

[37] E. M. Vedula, J. L. Alonso, M. A. Arnaout, J. L. Charest, PLOS ONE 2017, 12, e0184330.

[38] K. L. Sellgren, B. T. Hawkins, S. Grego, Biomicrofluidics 2015, 9, 061102.

[39] D. D. Nalayanda, Q. Wang, W. B. Fulton, T.-H. Wang, F. Abdullah, Journal of Pediatric Surgery 2010, 45, 45.

[40] K. Sharma, N. Dhar, V. V. Thacker, T. M. Simonet, F. Signorino-Gelo, G. W. Knott, J. D. McKinney, eLife 2021, 10, e66481.

[41] EMBO reports 2021, 22, e52744.

[42] B. D. Gastfriend, S. P. Palecek, E. V. Shusta, Current Opinion in Biomedical Engineering 2018, 5, 6.

[43] H. K. Wilson, S. G. Canfield, M. K. Hjortness, S. P. Palecek, E. V. Shusta, Fluids and Barriers of the CNS 2015, 12, 13.

[44] M. C. McCloskey, P. Kasap, S. D. Ahmad, S.-H. Su, K. Chen, M. Mansouri, N. Ramesh, H. Nishihara, Y. Belyaev, V. V. Abhyankar, et al., 2022, doi.org/10.1101/2022.03.28.486095.

[45] J. C. McDonald, D. C. Duffy, J. R. Anderson, D. T. Chiu, H. Wu, O. J. A. Schueller, G. M. Whitesides, ELECTROPHORESIS 2000, 21, 27.

[46] Ahmed, I. M. Joshi, S. Larson, M. Mansouri, S. Gholizadeh, Z. Allahyari, F. Forouzandeh, D. A. Borkholder, T. R. Gaborski, V. V. Abhyankar, Advanced Materials Technologies 2021, 6, 2001186.

[47] M. J. Williams, N. K. Lee, J. A. Mylott, N. Mazzola, A. Ahmed, V. V. Abhyankar, Micromachines 2019, 10, 360.

[48] V. V. Abhyankar, M. Wu, C.-Y. Koh, A. V. Hatch, PLOS ONE 2016, 11, e0156341.

[49] M. R. Hasan, S. S. S. Peri, V. P. Sabane, N. Mansur, J. X. Gao, K. T. Nguyen, J. A. Weidanz, S. M. Iqbal, V. V. Abhyankar, Biomed. Phys. Eng. Express 2018, 4, 025015.

[50] Ahmed, I. M. Joshi, M. Mansouri, A. M. Byerley, S. W. Day, T. R. Gaborski, V. V. Abhyankar, 2022, doi.org/10.1101/2022.02.04.479166.

[51] T. Salminen, J. Tithof, Y. Izhiman, E. A. Masters, M. C. McCloskey, T. R. Gaborski, D. H. Kelley, A. P. Pietropaoli, R. E. Waugh, J. L. McGrath, Integrative Biology 2020, 12, 275.

[52] P. Jaccard, New Phytologist 1912, 11, 37.

[53] Ivetic, Cell Tissue Res 2018, 371, 437.

[54] U. H. Von Andrian, P. Hansell, J. D. Chambers, E. M. Berger, I. Torres Filho, E. C. Butcher, K. E. Arfors, American Journal of Physiology-Heart and Circulatory Physiology 1992, 263, H1034.

[55] M. J. Mitchell, K. S. Lin, M. R. King, Biophysical Journal 2014, 106, 2243.

[56] J. S. Bredfeldt, Y. Liu, M. W. Conklin, P. J. Keely, T. R. Mackie, K. W. Eliceiri, J Pathol Inform 2014, 5, 28.

[57] F. Zittrell, CircHist, 2022.

[58] N. Wettschureck, B. Strilic, S. Offermanns, Physiological Reviews 2019, 99, 1467.

[59] F. Dewey Jr., S. R. Bussolari, M. A. Gimbrone Jr., P. F. Davies, Journal of Biomechanical Engineering 1981, 103, 177.

[60] N. E. Ashpole, D. R. Overby, C. R. Ethier, W. D. Stamer, Investigative Ophthalmology & Visual Science 2014, 55, 8067.

[61] F. Moazzam, F. A. DeLano, B. W. Zweifach, G. W. Schmid-Schönbein, Proceedings of the National Academy of Sciences 1997, 94, 5338.

[62] S. Fukuda, G. W. Schmid-Schönbein, Proceedings of the National Academy of Sciences 2003, 100, 13152.

[63] Ahmed, I. M. Joshi, M. Mansouri, N. N. N. Ahamed, M.-C. Hsu, T. R. Gaborski, V. V. Abhyankar, American Journal of Physiology-Cell Physiology 2021, 320, C1112.

[64] M. J. Jensen, G. Goranovi, H. Bruus, J. Micromech. Microeng. 2004, 14, 876.

[65] M.-C. Hsu, M. Mansouri, N. N. N. Ahamed, I. M. Joshi, A. Ahmed, D. A. Borkholder, V. V. Abhyankar, 2022, doi.org/10.1101/2022.04.03.486540.

[66] M. W. Toepke, D. J. Beebe, Lab Chip 2006, 6, 1484.

[67] G. Vunjak-Novakovic, K. Ronaldson-Bouchard, M. Radisic, Cell 2021, 184, 4597.

