## Supporting Information for "The Modular μSiM Reconfigured: Integration of Microfluidic Capabilities to Study in vitro Barrier Tissue Models under Flow"

### S1. Molds for the fabrication of flow module and seeding stencil

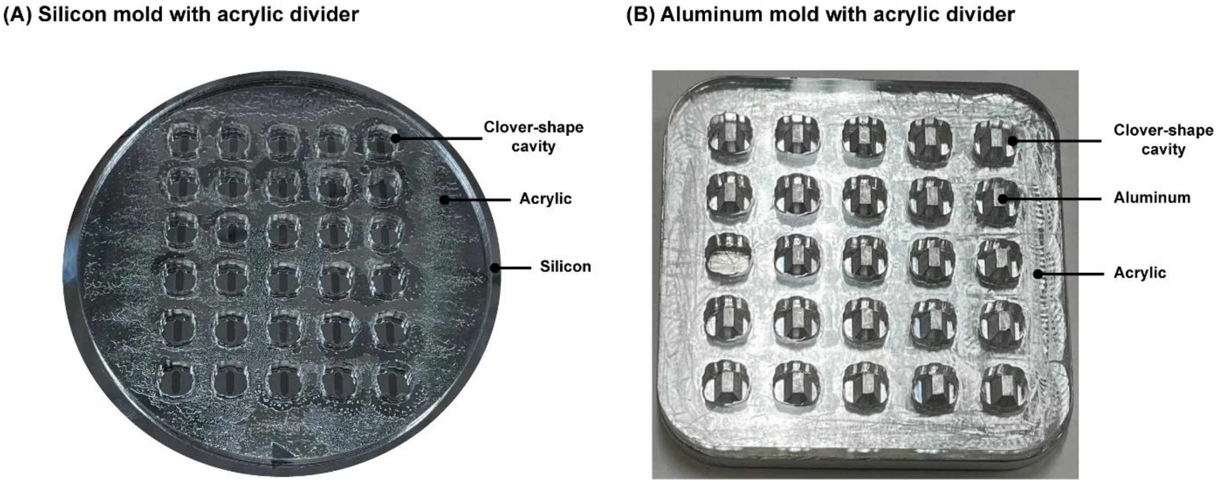


**Figure S1.** (A) Silicon mold with microchannel features bonded to an acrylic divider for fabricating clover-shape flow modules. (B) Machined aluminum mold with tapered features bonded to an acrylic divider for fabricating clover-shape seeding stencils. Both dividers comprise cavities that confines PDMS to get a clover-shape during curing on the hot plate.

### S2. Flow circuit

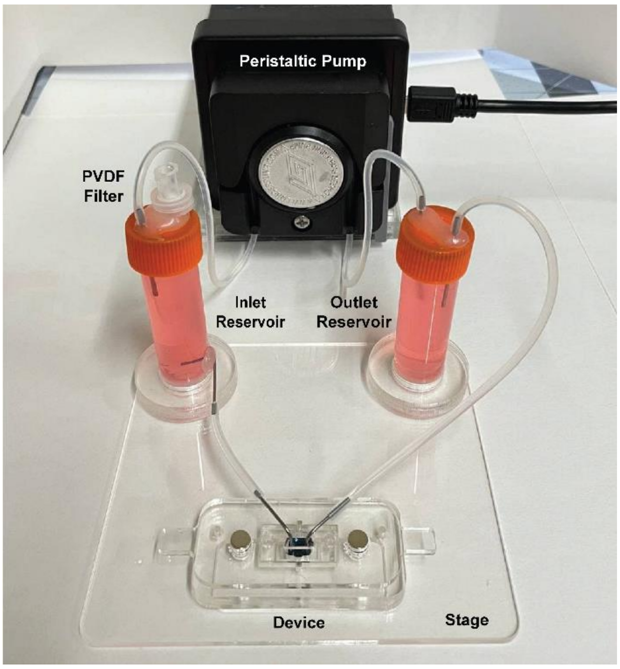


**Figure S2.** Custom-made flow circuit for experiments with continuous fluid flow. The setup includes a peristaltic pump for media circulation throughout the system, two reservoirs for supplying cell media and damping flow fluctuations, and an acrylic stage to hold the components in place. The PVDF filter is used to equilibrate cell media with environmental conditions in the incubator.

### S3. Particle image velocimetry (PIV)

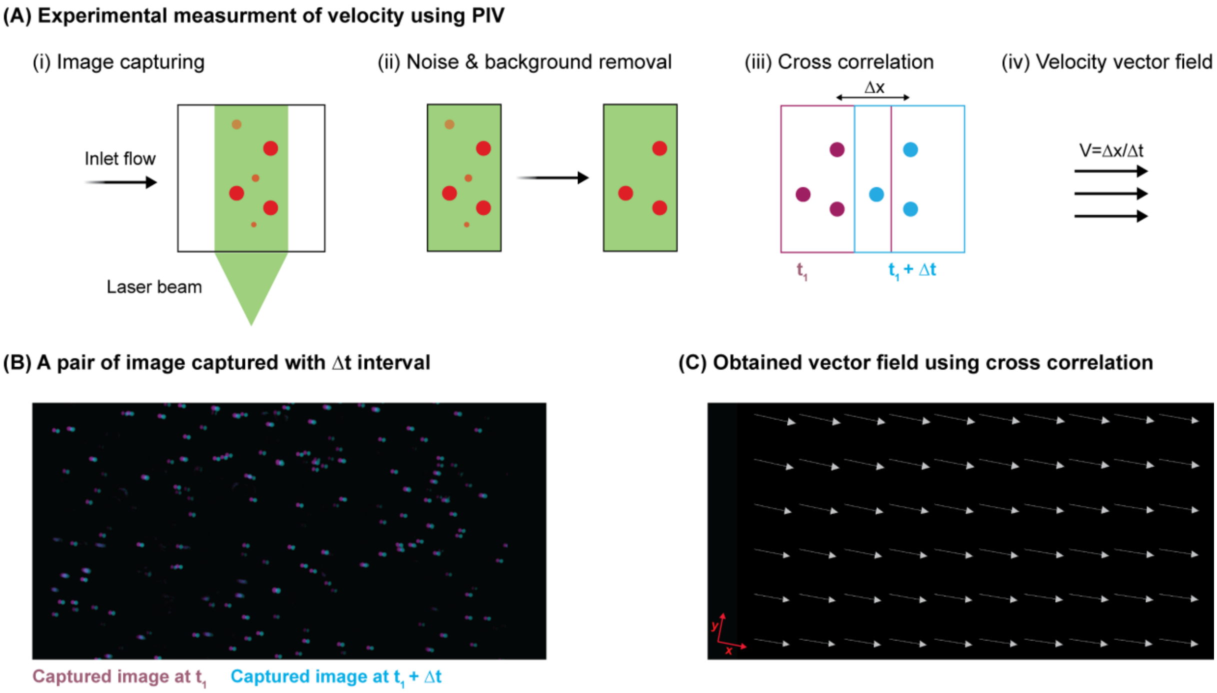


**Figure S3.** Schematic illustration of the experimental flow analysis using particle image velocimetry technique. (A) (i) Laser beam excites fluorescent beads in the region of interest and several pairs of images are captured. (ii) Noise and out-of-focus beads are removed from each image using image processing algorithms. (iii) Cross-correlation is applied to images of beads to find displacement between the two paired images. (iv) Velocity vectors are calculated on a structured grid for each pair of images and the mean velocity among all pairs is demonstrated as the velocity vector field. (B) A pair of image captured with a time interval of 100 µs. (C) Generated vector field using average velocity obtained from 50 pair of images.

### S4. Geometry and dimensions of the flow module

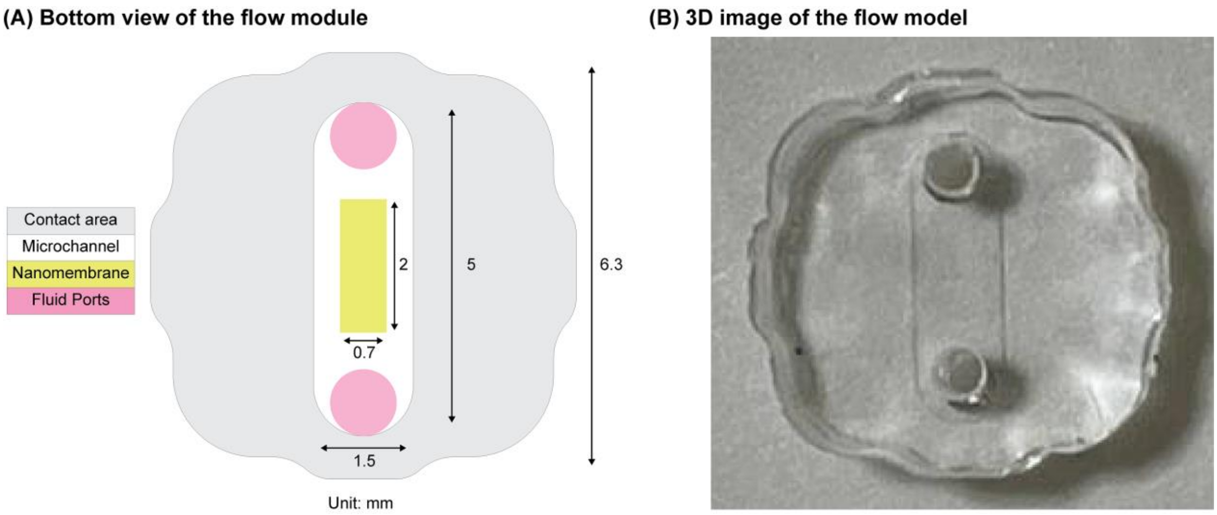


**Figure S4.** (A) Schematic illustration of the contacting interface between m-µSiM membrane chip and the flow module. The fluid flow enters and exits the microchannel from fluid ports shown in pink. (B) 3D image of the PDMS flow module. Both m-µSiM well and the flow module have a clover-shape to promote self-alignment. All dimensions are in mm.

### S5. Neutrophil introduction and deposition onto the membrane

As a particle moves through the microchannel along with the flow, it spends a certain amount of time over the membrane (**Figure S5**). This parameter is called "residence time" ($t_f$) and can be calculated by equation (S1) where L is the length of the membrane and $V_f$ is the velocity of the fluid at the given height of the particle with respect to the membrane. There is another time scale called "settling time" ($t_s$) that describes the time it takes for a particle to settle from a position h onto the membrane surface which can be calculated by equation (S2). H is the height of the particle with respect to the membrane and Vs is the settling velocity of the particle which can be calculated using Stokes' law as described by equation (S3).

Residence time: $t_f = \frac{L}{V_f}$ (S1)

Settling time: $t_s = \frac{H}{V_s}$ (S2)

Settling velocity: $V_s = \frac{gd^2(\rho_p - \rho_m)}{18\mu}$ (S3)

Parameters $d, \rho_p, \rho_m, \mu$ represent particle diameter, particle density, medium density, and medium viscosity, respectively.

In a situation where the settling time is less than the residence time ($t_s < t_f$), the particle can settle onto the membrane before it is swept past the membrane. To compare settling and residence times of particles at different heights, we obtained $V_f$ from the COMSOL simulation and calculated Vs using Stokes' Law for different flow rates. Assuming that neutrophils are distributed uniformly in the solution, our calculations showed that an inlet flow rate of 10 $\mu L.min^{-1}$ results in the deposition of 20 % of neutrophil onto the membrane, which provides an adequate number of neutrophils on the membrane for transmigration study. This is an acceptable flow rate since it prevents neutrophil activation while providing adequate neutrophil deposition. Higher flow rates lead to neutrophil activation and lower flow rates result in the settlement of the majority of neutrophils in the tubing. If more neutrophils are desired on the membrane surface, we suggest increasing the neutrophil concentration in the initial solution or increasing the time the flow is maintained (introduction time).

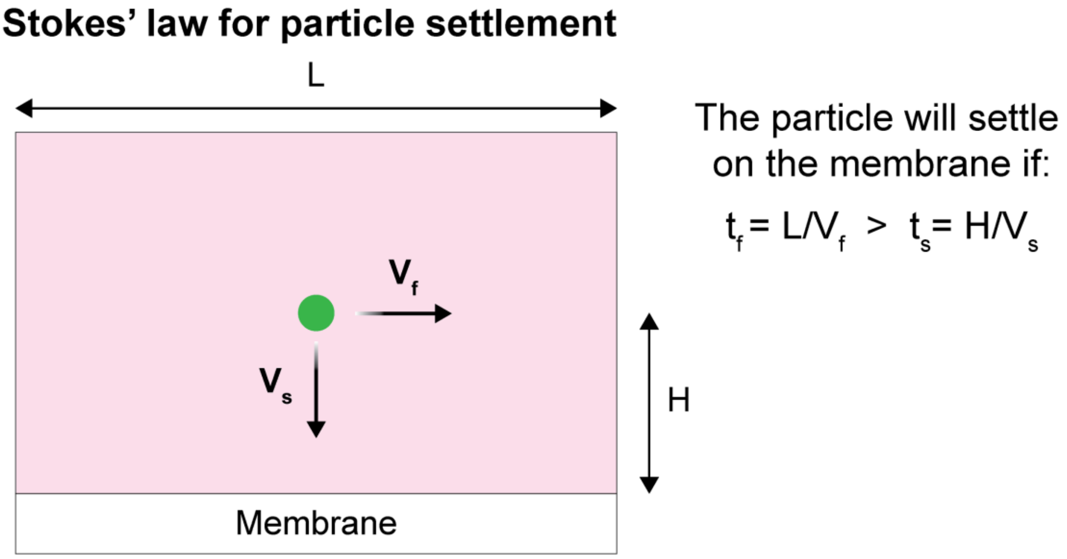


**Figure S5.** Schematic demonstration of Stokes' law for a spherical particle that follows the fluid velocity. When the fluid is incompressible and the flow is fully developed, Stokes' law provides a mathematical model for calculating the settling velocity of the particle and approximating the required time for the particle to settle on the surface, $t_s$. We compare this to the time required to move through the flow module, $t_f$, as an estimate of the likelihood of a particle settling onto the membrane.

#### S6. Quantification of neutrophil activation using flow cytometry

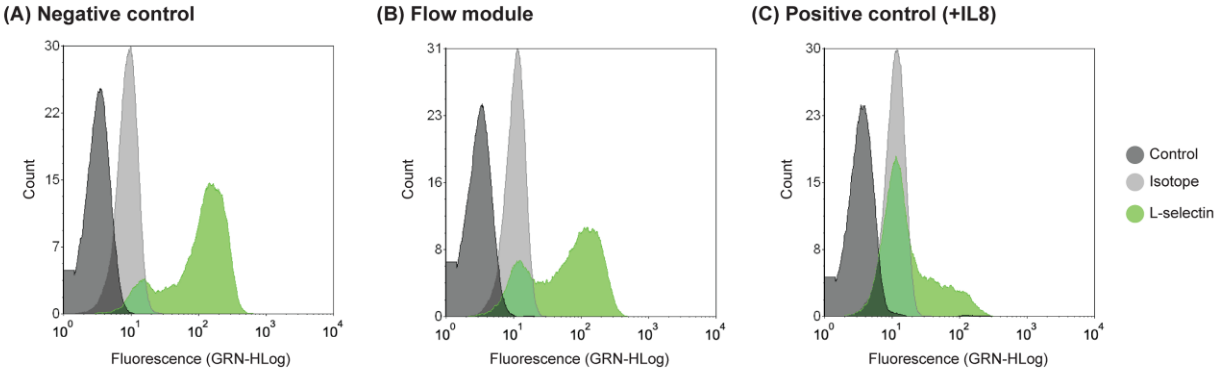


**Figure S6.** Flow cytometry analysis of neutrophils using L-selectin as an activation biomarker. (A) Post isolation stock as the negative control, (B) neutrophils flowed through the device showing a similar trend to the negative control, (C) IL8 treated neutrophils as the positive control showing loss of L-selectin due to activation.

### S7. Mimicking tissue side in the bottom channel of m-μSiM by incorporating collagen I

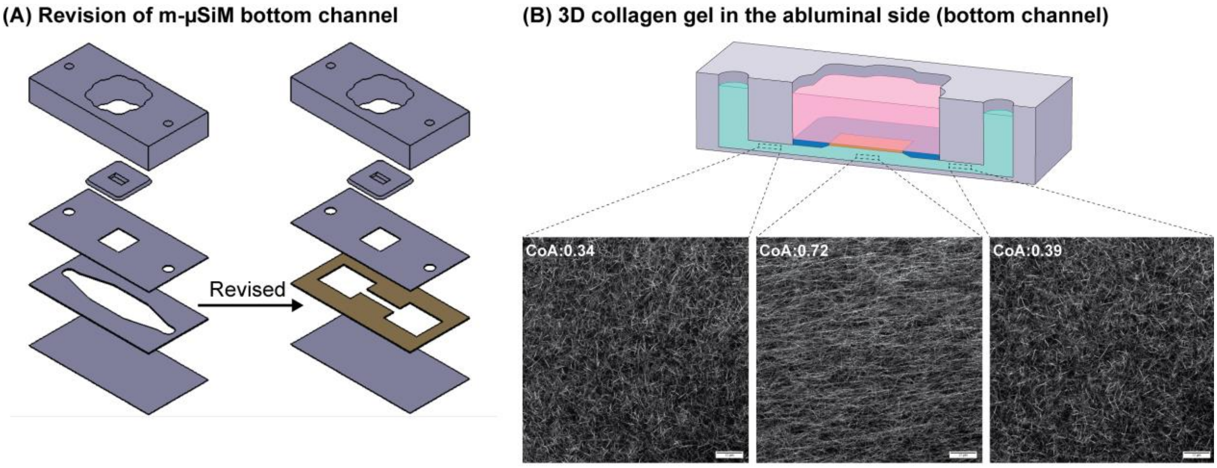


**Figure S7.** Customization of the m-μSiM component to incorporate 3D gel with aligned collagen fibers below the membrane. (A) The bottom channel of the m-μSiM is revised into a segmented channel to align collagen fibers based on extensional strain. (B) Demonstration of fibers at three different locations across the bottom channel. CoA represents the coefficient of fiber alignment (COA=0: complete randomness, COA=1: complete alignment).

### S8. Characteristics of the fluid flow in the flow module

**Table S1.** Characteristics of fluid flow within the flow module at different flow rates

| Inlet flow rate (μl.min$^{-1}$) | Average V at midline of the flow path-PIV (mm.s$^{-1}$) | Average V at midline of the flow path-COMSOL (mm.s$^{-1}$) | Shear stress on the membrane surface (dynes.cm$^{-2}$) | Maximum shear stress within the flow path (dynes.cm$^{-2}$) |
|---|---|---|---|---|
| 10 | 0.89 ± 0.07 | 0.83 | 0.18 | 0.40 |
| 100 | 8.73 ± 0.1 | 8.31 | 1.85 | 3.93 |
| 200 | 15.01 ± 1.09 | 16.62 | 3.69 | 7.96 |
| 500 | 40.39 ± 0.84 | 41.55 | 9.23 | 20.89 |
| 1000 | 86.57 ± 4.49 | 83.10 | 18.45 | 42.49 |
